# APOBEC3G expression marks a TMB-high, T cell–inflamed tumor state and is associated with response to immune checkpoint blockade in multiple cancer cohorts

**DOI:** 10.64898/2026.07.08.737369

**Authors:** Kelly E. Butler, Dhanusha Yesudhas, Bilal Lone, A. Rouf Banday

## Abstract

Immune checkpoint therapies have transformed clinical practice; however, reliable biomarkers to predict response remain limited. Tumor mutational burden (TMB) has emerged as an important biomarker because it is thought to reflect neoantigen load, yet its predictive utility has been inconsistent. This limitation may partly arise because TMB primarily captures tumor-intrinsic immunogenicity, which is heterogeneous and does not fully reflect the state of antitumor immunity. To identify transcriptomic surrogates that capture both high mutational burden and antitumor immune activation, we investigated whether mRNA expression of mutagenic APOBEC3 family members could serve as surrogates for high TMB and T cell–rich tumors. Using a pan-cancer computational framework, we evaluated the association of four APOBEC3 genes with mutational burden, neoantigen load, immune infiltration, and immune checkpoint blockade response. Among *APOBEC3A, APOBEC3B, APOBEC3G*, and *APOBEC3H, APOBEC3G* emerged as the strongest and most consistent marker of a TMB^high^CD8^high^ and Neoantigen^high^CD8^high^ tumor phenotypes. Single-cell analyses further demonstrated that *APOBEC3G* is enriched in both malignant cells and T cells compared with other *APOBEC3* family members, with *APOBEC3G*-positive CD8^+^ T cells exhibiting elevated activation markers including GZMB and IFNG. Importantly, retrospective analyses of 50 immune checkpoint blockade cohorts showed that *APOBEC3G* had the most consistent association among *APOBEC3* family members with treatment response and clinical outcomes. Together, these findings identify *APOBEC3G* as a candidate transcriptomic marker of a TMB-associated, T cell–inflamed tumor state linked to immune-checkpoint blockade benefit, warranting further prospective validation.

## Introduction

Immune checkpoint therapies have transformed oncology; however, robust and broadly applicable biomarkers of response remain elusive. Tumor mutational burden (TMB) has been widely adopted as a surrogate for neoantigen load, yet its predictive performance is inconsistent across tumor types and clinical settings (1, 2). This limitation may reflect that TMB captures tumor-intrinsic mutational processes without fully accounting for the immune context required for effective anti-tumor responses (3). The APOBEC3 family of cytosine deaminases represents a major endogenous source of somatic mutagenesis in cancer, predominantly driving C>T and C>G substitutions in specific sequence contexts (4, 5). While APOBEC3A and APOBEC3B are most strongly linked with SBS2 and SBS13 mutational signatures, APOBEC3G has been associated with a distinct mutational signature (5, 6). Certain APOBEC3H haplotypes have also been reported to exhibit mutagenic activity (7). More broadly, APOBEC3 activity contributes to diffuse hypermutation, kataegis, and chromosomal instability (8, 9). In bladder, breast, cervical, head and neck, and lung malignancies, APOBEC3-induced mutations can account for a substantial fraction of overall tumor mutational burden, reaching over two-thirds in some cases. (10). Notably, *APOBEC3* expression has been linked to both mutational load and tumor immune activity (11-13), raising the possibility that *APOBEC3* transcripts may serve as integrated surrogates of tumor mutagenicity and immune activation.

## Results and Discussion

We therefore hypothesized that *APOBEC3* gene expression could serve as a composite, transcript-based biomarker of tumor immunogenicity. To address this, we first defined TMB^high^CD8^high^ and Neoantigen^high^CD8^high^ tumor phenotypes across 32 cancer types using residual-based regression analyses (**Supplementary methods**), then evaluated associations between these phenotypes and expression of mutagenic APOBEC3 family members. Although expression of all four mutagenic *APOBEC3s* was broadly associated with the TMB^high^CD8^high^ and Neoantigen^high^CD8^high^ tumor phenotypes across multiple cancer types, *APOBEC3G* exhibited the strongest and most consistent associations (**Figure 1a, b; Supplementary Figures 1 and 2**). Pooled Cochran–Mantel–Haenszel analyses across multiple CD8 readouts further confirmed that *APOBEC3G* showed the strongest enrichment for tumors characterized by both the TMB^high^CD8^high^ and Neoantigen^high^CD8^high^ phenotypes, with *APOBEC3H* showing the next most consistent association (**Supplementary Figure 3**). Complementary cancer-type-adjusted regression analyses supported these findings and showed that *APOBEC3G* remained associated with composite TMB^high^CD8^high^ and Neoantigen^high^CD8^high^ states after adjustment for tumor purity (**Supplementary Figure 4a, b**).

**Figure 1.**
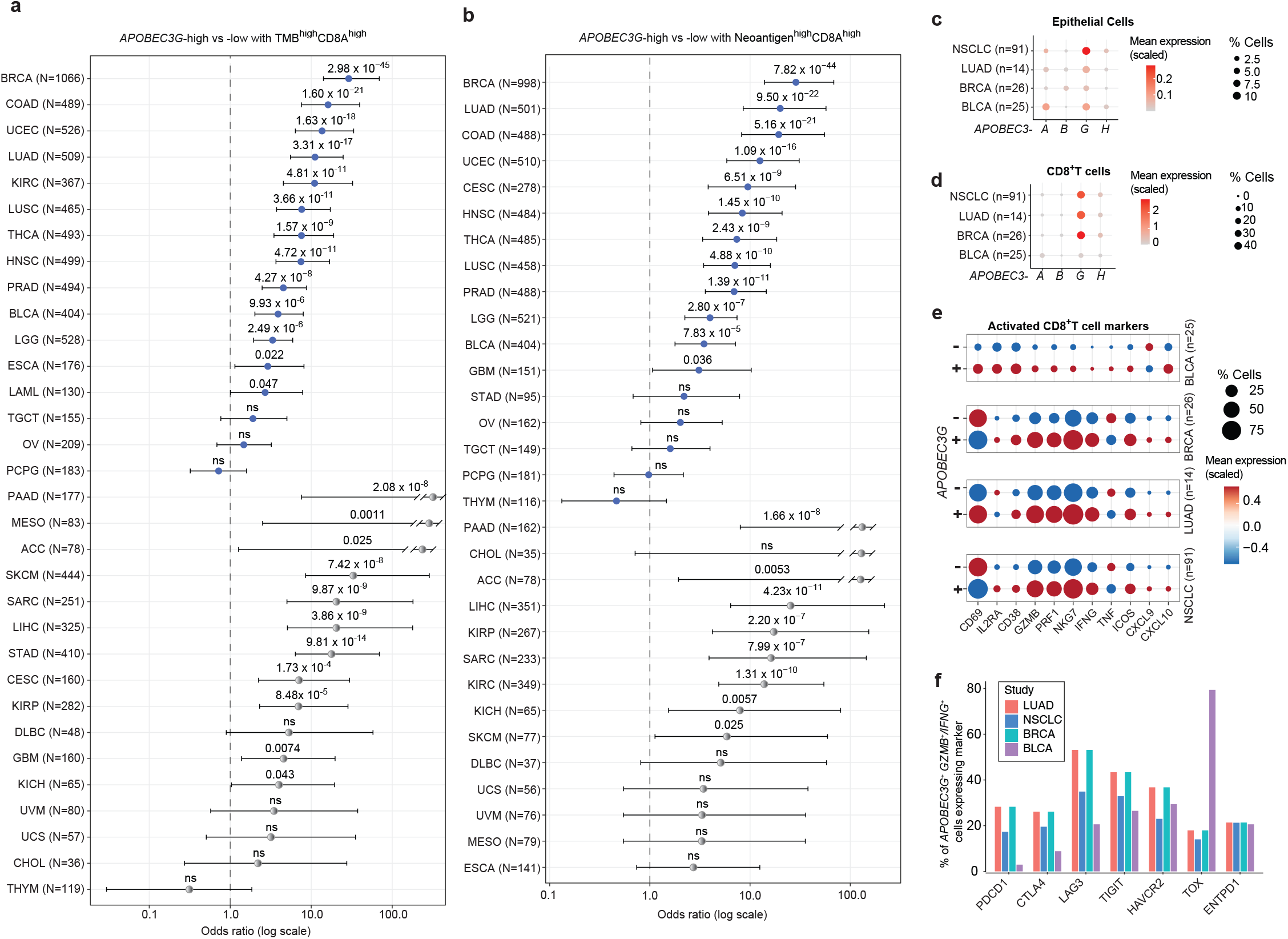
*APOBEC3G* is the strongest *APOBEC3* family correlate of TMB^high^CD8^high^ phenotype and is enriched in tumor and T cell compartments. **(a)** Forest plot showing odds ratios (ORs) and 95% confidence intervals (CIs) for the association between *APOBEC3G* expression (high vs. low) and a TMB^high^CD8^high^ phenotype across cancer types using *CD8A* as the CD8 readout. The full names of the abbreviated cancers, according to TCGA nomenclature, are provided in the Supplementary Materials (methods section). Cancer types are ordered by effect size. Gray points indicate estimates derived from sparse data (any contingency table cell <5), which were retained but interpreted with caution. **(b)** Same analysis as in (**a**), evaluating neoantigen load in place of TMB. *APOBEC3G* shows consistent enrichment for tumors with Neoantigen^high^CD8^high^ phenotype across multiple cancer types. **(c)** Single-cell RNA sequencing analysis showing mean expression (color scale) and percentage of expressing cells (dot size) for mutagenic APOBEC3 family members (*APOBEC3A, APOBEC3B, APOBEC3G, APOBEC3H*) in malignant epithelial cells across tumor types. **(d)** Single-cell analysis of *APOBEC3* expression in CD8^+^ T cells across tumor types, showing mean expression and percentage of expressing cells. **(e)** Expression of activation markers in CD8^+^ T cells stratified by *APOBEC3G* expression. *APOBEC3G*-positive CD8^+^ T cells exhibit increased expression of activation markers (e.g., *GZMB* and *IFNG*), consistent with an activated, antigen-experienced phenotype. **(f)** Percentage of *APOBEC3G*-positive cells co-expressing immune checkpoint and activation markers across immunotherapy-relevant genes, stratified by study and cancer type.

Single-cell analyses provided additional biological insight. Because APOBEC3-family activity is most relevant in tumor types with prominent APOBEC3-medaited mutational processes, we first examined representative datasets from APOBEC3-mutagenesis–associated cancers, including non-small cell lung cancer (NSCLC), lung adenocarcinoma (LUAD), breast cancer (BRCA), and bladder cancer (BLCA) (**Figure 1c, d**). Among the APOBEC3 family members examined, *APOBEC3G* showed the most consistent enrichment in both malignant epithelial cells and CD8^+^ T cells across these datasets. To assess whether this pattern extended beyond these datasets, we analyzed additional independent single-cell cohorts, including newly generated bladder cancer data and public kidney cancer, liver cancer, and melanoma cohorts (**Supplementary Figure 5a, b**). These analyses similarly supported *APOBEC3G* enrichment in tumor and immune compartments. Within CD8^+^ T cells, *APOBEC3G*-positive cells showed higher expression of activation and effector markers, including *GZMB* and *IFNG*, compared with *APOBEC3G*-negative CD8^+^T cells (**Figure 1e and Supplementary Figure 5c**). In addition, a substantial fraction of APOBEC3G^+^GZMB^+^IFNG^+^ CD8+ T cells co-expressed inhibitory or antigen-experience markers, including *PDCD1, TIGIT, HAVCR2, LAG3, CTLA4, ENTPD1*, and *TOX* (**Figure 1f and Supplementary Figure 5d**). These findings suggest that *APOBEC3G* expression reflects a composite tumor–immune state, with contributions from both malignant and immune-cell compartments. In bulk data from TCGA tumors, *APOBEC3G* expression was also positively associated with multiple CD8/T-cell-inflamed markers in cancer-type-adjusted models, including analyses additionally accounting for tumor purity (**Supplementary Figure 4c**). These analyses further support the interpretation that *APOBEC3G* marks a T cell–inflamed tumor state rather than mutational burden alone.

To assess clinical relevance, we evaluated the expression of *APOBEC3* family genes across 50 immune-checkpoint blockade cohorts spanning multiple cancer types (>5,000 patients). Among the *APOBEC3* family members evaluated, *APOBEC3G* showed the most consistent association with immune-checkpoint blockade response rates and survival outcomes across retrospective cohorts (**Figure 2a-c**). Associations for *APOBEC3A, APOBEC3B*, and *APOBEC3H* were observed in fewer endpoints and cohorts (**Supplementary Figure 6)**.

**Figure 2.**
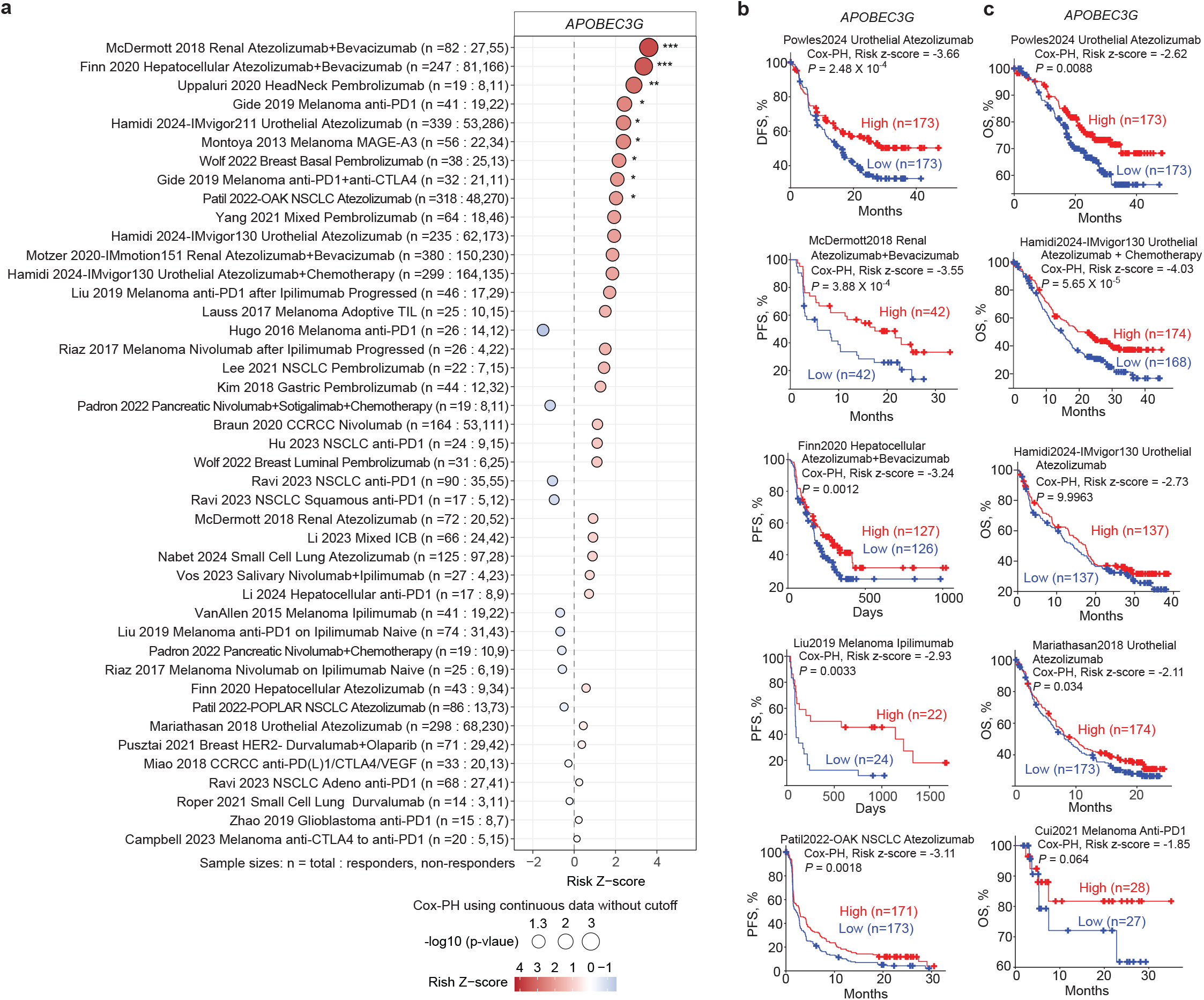
*APOBEC3G* mRNA expression is associated with clinical immunotherapy response and outcomes across independent cancer cohorts. **(a)** Dot plot summarizing associations between *APOBEC3G* expression and clinical response (responders vs. non-responders) across multiple immunotherapy-treated cohorts. Clinical and molecular data were obtained and curated from the Cancer Immunology Data Engine (CIDE). Signed Wald z-statistics and corresponding P values were obtained from logistic regression models fitted using continuous expression values without predefined cutoffs. Colors indicate the direction and magnitude of association for higher *APOBEC3G* expression (red, favorable association; blue, unfavorable association). Circle size reflects statistical significance (‒log10 *P* value), and significance levels are denoted by asterisks (*P* < 0.05, *P* < 0.01, *P* < 0.001). Sample sizes are shown as total patients, followed by responders and non-responders. **(b, c)** Representative Kaplan–Meier curves illustrating associations between *APOBEC3G* expression and progression-free survival (PFS), disease-free survival (DFS), or overall survival (OS) in cohorts that showed the strongest associations. Risk z-statistics and corresponding *P* values were derived from Cox proportional hazards models.

Collectively, these data support a model in which bulk *APOBEC3G* expression marks a composite tumor–immune state in which mutational burden and antitumor immune activation coexist, particularly in cancers with high mutational activity. Unlike TMB, which reflects mutational load without directly measuring immune contexture, *APOBEC3G* expression appears to mark tumors in which mutational burden coexists with a T cell–inflamed microenvironment, a state associated with clinical benefit from immune-checkpoint blockade. These findings are based on retrospective transcriptomic analyses of independent cohorts with substantial heterogeneity in tumor type, treatment regimen, sequencing platform, preprocessing workflow, response definition, and sample size; thus, biomarker performance estimates should be interpreted cautiously and require prospective validation. A further limitation of bulk transcriptomic analyses is that *APOBEC3G* expression cannot be definitively assigned to malignant-cell-intrinsic versus immune-cell-derived sources. Therefore, we do not interpret bulk *APOBEC3G* as a single-compartment marker. Rather, our data support *APOBEC3G* as a candidate composite transcriptomic biomarker that captures both tumor-associated APOBEC3 biology and immune-cell-associated inflammation within the tumor microenvironment.

Importantly, *APOBEC3G* is not intended to replace TMB as an established biomarker, but rather to complement mutation-based metrics by capturing features of both mutational burden and immune activation in a single transcript-based measure. *APOBEC3G* expression may offer practical value in research or clinical contexts where tumor RNA-seq, targeted expression panels or transcriptomic profiling are already available; however, prospective validation will be required before clinical implementation. In summary, *APOBEC3G* emerged as the most consistent *APOBEC3* family marker of immune-checkpoint blockade benefit across retrospective cancer cohorts. *APOBEC3G* thus warrants prospective evaluation as a candidate transcriptomic marker of a TMB-associated, T cell–inflamed tumor state linked to immune-checkpoint blockade response. More broadly, our findings suggest that transcript-based biomarkers integrating tumor-intrinsic mutational processes with immune context may complement mutation-based metrics and improve biomarker development for immune checkpoint blockade.

## Materials and Methods

### Human Tumor Omics Datasets and Data Sources

TCGA pan-cancer RNA-sequencing, mutation, neoantigen, immune-infiltration, and clinical annotation data were accessed through the UCSC Xena Browser, cBioPortal, and previously curated pan-cancer immunogenomic resources. TCGA cancer types were included when *APOBEC3* family expression estimates, mutational-burden or neoantigen-burden estimates, and CD8/T-cell infiltration readouts were available. No disease-specific inclusion or exclusion criteria were applied beyond availability of the required molecular and clinical annotations.

For single-cell analyses, we analyzed whole-tumor single-cell or single-nucleus RNA-sequencing datasets from 25 bladder tumors (GEO: GSE169379), 26 breast tumors (GEO: GSE176078), 14 lung adenocarcinomas (GEO: GSE123902), and a non–small cell lung cancer dataset comprising 91 patients (GEO: GSE154826). We additionally analyzed whole-tumor single-cell RNA-sequencing datasets from 5 bladder tumors generated by our group (GEO: GSE319003), 10 melanoma tumors (GEO: GSE148190), 15 liver tumors (GEO: GSE125449), and 6 kidney tumors from the Broad Single Cell Portal dataset SCP1288, with raw data deposited in dbGaP under accession phs002065.v1.p1.

Immune checkpoint blockade cohorts were obtained from the Cancer Immunology Data Engine (CIDE). We downloaded and analyzed the available processed cohort-level transcriptomic matrices and matched clinical annotation from CIDE for cohorts with pretreatment transcriptomic data and response and/or survival outcomes. Cohorts were included when *APOBEC3* family expression could be quantified from the available transcriptomic profiles. These datasets represented 43 studies comprising 50 treatment cohorts and 5,363 patients with at least one available clinical endpoint. Endpoint-specific sample sizes varied across cohorts according to the availability of response, RECIST, progression-free survival, or overall survival annotation.

Additional genomics datasets used in this study are referenced and described in the relevant sections below. The full names of the abbreviated cancers are: Adrenocortical Carcinoma (ACC), Bladder Urothelial Carcinoma (BLCA), Breast Invasive Carcinoma (BRCA), Cervical Squamous Cell Carcinoma and Endocervical Adenocarcinoma (CESC), Cholangiocarcinoma (CHOL), Colon Adenocarcinoma (COAD), Lymphoid Neoplasm Diffuse Large B-cell Lymphoma (DLBC), Esophageal Carcinoma (ESCA), Glioblastoma Multiforme (GBM), Head and Neck Squamous Cell Carcinoma (HNSC), Kidney Chromophobe (KICH), Kidney Renal Clear Cell Carcinoma (KIRC), Kidney Renal Papillary Cell Carcinoma (KIRP), Acute Myeloid Leukemia (LAML), Brain Lower Grade Glioma (LGG), Liver Hepatocellular Carcinoma (LIHC), Lung Adenocarcinoma (LUAD), Lung Squamous Cell Carcinoma (LUSC), Mesothelioma (MESO), Ovarian Serous Cystadenocarcinoma (OV), Pancreatic Adenocarcinoma (PAAD), Pheochromocytoma and Paraganglioma (PCPG), Prostate Adenocarcinoma (PRAD), Rectum Adenocarcinoma (READ), Sarcoma (SARC), Skin Cutaneous Melanoma (SKCM), Stomach Adenocarcinoma (STAD), Testicular Germ Cell Tumors (TGCT), Thyroid Carcinoma (THCA), Thymoma (THYM), Uterine Corpus Endometrial Carcinoma (UCEC), Uterine Carcinosarcoma (UCS), Uveal Melanoma (UVM).

### Single-cell and single-nucleus RNA-sequencing analysis

Single-cell and single-nucleus RNA-sequencing datasets were analyzed as whole-tumor datasets to evaluate the cellular compartments contributing to APOBEC3 family expression. We prioritized datasets that included tumor and immune compartments rather than datasets generated exclusively from sorted malignant or immune-cell populations. Cell-type annotations were used as provided by the original studies when available; otherwise, major cellular compartments were assigned using canonical lineage markers. *APOBEC3* family expression was summarized by cell type using the percentage of expressing cells and average expression. For CD8^+^ T-cell analyses, *APOBEC3G*-positive and *APOBEC3G*-negative CD8^+^ T cells were compared for expression of activation and effector markers, including *GZMB, IFNG*, and immune checkpoint-associated genes.

### Residual-based identification of TMB^high^CD8^high^ and Neoantigen^high^CD8^high^ tumors

Tumors were analyzed separately within each cancer type. CD8^+^ T-cell abundance was quantified using five independent readouts: CD8A expression, CD8B expression, xCell CD8^+^ T-cell score (14), CIBERSORT absolute CD8^+^ T-cell score (15), and TIMER CD8^+^ T-cell score (16). Tumor mutational burden (TMB) was represented by nonsynonymous mutation rate (Nonsilent_Mutation_Rate), and neoantigen load was represented by Total_Neoantigens as previously computed (17). For each cancer type and CD8 readout, linear regression models were fitted:

CD8 readout ∼ TMB, and CD8 readout ∼ Total Neoantigens

Residuals were extracted for each sample. A positive residual indicates that the observed CD8 signal exceeds that expected based on TMB or neoantigen load within that cancer type. Within each cancer type, samples were classified as TMB-high or neoantigen-high if their values were greater than or equal to the cancer-specific median of Nonsilent_Mutation_Rate or Total_Neoantigens, respectively. Samples were classified as CD8-high if the corresponding CD8 readout exceeded the cancer-specific median. A sample was defined as belonging to the TMB^high^CD8^high^ residual-enriched group if it satisfied all of the following criteria: a, TMB ≥ cancer-specific median; b, CD8 readout ≥ cancer-specific median; and c, Positive residual from the CD8 ∼ TMB regression. Similarly, a sample was assigned to the Neoantigen^high^CD8^high^ residual-enriched group. This framework was applied independently for each of the five CD8 readouts.

### Association of APOBEC3 expression with residual-enriched immunogenomic groups

Expression of *APOBEC3A, APOBEC3B, APOBEC3G*, and *APOBEC3H* was analyzed separately. Expression values were analyzed on a log2(TPM + 1) scale. Within each cancer type, samples were dichotomized into APOBEC-high and APOBEC-low groups using the cancer-specific median expression of each gene. For each cancer type, 2×2 contingency tables were constructed comparing APOBEC-high versus APOBEC-low status with membership in the residual-enriched groups defined above. Analyses were performed separately for: a) TMB^high^CD8^high^ residual-enriched tumors, and b) Neoantigen^high^CD8^high^ residual-enriched tumors and repeated across all five CD8 readouts. To address zero counts and ensure stable estimation, a Haldane–Anscombe continuity correction (+0.5 to all cells) was applied prior to statistical testing. Odds ratios (ORs), 95% confidence intervals (CIs), and P values were computed using Fisher’s exact test. Odds ratios greater than 1 indicate enrichment of APOBEC-high tumors within the residual-enriched (double-high) group. Estimates were classified as sparse when any cell in the original (uncorrected) contingency table contained fewer than five observations. These estimates were retained to preserve completeness but were flagged and interpreted with caution due to increased statistical uncertainty.

### Cochran–Mantel–Haenszel analysis across cancer types

To obtain pan-cancer summary estimates while accounting for tumor-type stratification, associations between APOBEC-high status and residual-enriched group membership were evaluated using the Cochran–Mantel–Haenszel (CMH) test. For each APOBEC3 gene, CD8 readout, and immunogenomic grouping (TMB or neoantigen), cancer-specific 2×2 contingency tables were combined into a stratified 2×2×K table (K = number of cancer types). The CMH test was used to estimate common odds ratios, 95% confidence intervals, and two-sided P values across strata. This analysis tested whether tumors with high APOBEC3 expression had increased odds of belonging to TMB^high^CD8^high^ or Neoantigen^high^CD8^high^ residual-enriched groups, while controlling for cancer type.

### Cancer-type- and tumor-purity-adjusted sensitivity analyses

To complement the stratified Cochran–Mantel–Haenszel analyses, we performed cancer-type-adjusted regression analyses evaluating the association between *APOBEC3* family expression and composite TMB^high^CD8^high^ or Neoantigen^high^CD8^high^ tumor states. For each *APOBEC3* gene, expression values were standardized, and logistic regression models were fitted with composite tumor-state membership as the outcome and *APOBEC3* expression as the predictor, adjusting for cancer type as a categorical covariate. Odds ratios are reported per one standard deviation increase in *APOBEC3* expression.

To evaluate whether *APOBEC3G* associations were explained by tumor purity, additional logistic regression models were fitted for *APOBEC3G* with cancer type and tumor purity included as covariates. Tumor purity estimates were obtained from available TCGA pan-cancer purity annotations. We also evaluated associations between *APOBEC3G* expression and immune/CD8-related markers using linear regression models adjusted for cancer type alone or for both cancer type and tumor purity. *APOBEC3G* expression and immune-marker values were standardized before modeling, and standardized beta coefficients with 95% confidence intervals were reported.

### Analysis of transcriptomic and clinical outcomes in immune checkpoint blockade cohorts

Processed pretreatment transcriptomic profiles and matched clinical outcome annotations for immune checkpoint blockade-treated cohorts were downloaded from the Cancer Immunology Data Engine (CIDE), accessed in December 2025 (18). Included cohorts had available APOBEC3 family expression estimates and at least one clinical endpoint, including objective response, RECIST response category, progression-free survival, or overall survival. Treatment cohorts primarily included anti-PD-1, anti-PD-L1, anti-CTLA-4, or combination checkpoint blockade regimens where available.

For each cohort, *APOBEC3A, APOBEC3B, APOBEC3G*, and *APOBEC3H* expression values were analyzed using the processed expression matrices provided by CIDE. Parallel analyses were performed for datasets also available through dbGaP for cross-validation. Associations between APOBEC3 expression and binary treatment response were evaluated using logistic regression. Associations with progression-free survival or overall survival were evaluated using Cox proportional hazards regression. Analyses were performed at the cohort level using available matched transcriptomic and clinical annotations; endpoint-specific sample sizes differed depending on the availability of response, RECIST, PFS, or OS data. For logistic and Cox regression models, APOBEC3 expression was modeled as a continuous variable within each cohort without predefined expression cutoffs. Statistical significance was assessed using Wald test P values.

### Reporting and statistics

For each *APOBEC3* gene, analyses were performed across all five CD8 readouts and for both TMB-based and neoantigen-based definitions. Results are reported as common odds ratios with 95% confidence intervals and two-sided Cochran–Mantel–Haenszel (CMH) P values. Cancer-specific sample counts were retained for all strata. For immunotherapy cohorts, analyses were performed using endpoint-specific sample sets; therefore, the number of samples included for response, RECIST, PFS, and OS analyses could differ within the same cohort. All statistical tests were two-sided, and no formal multiple testing correction was applied given the hypothesis-driven nature of the analyses; results were interpreted based on effect size, consistency across readouts, and nominal significance thresholds. Statistical analyses were performed using R.

## List of abbreviations

(APOBEC3): Apolipoprotein B mRNA-editing enzyme, catalytic polypeptide 3
(TMB): tumor mutation burden

## Declarations

### Ethics approval and consent to participate

Not applicable

### Availability of data and materials

The datasets analyzed in the current study are available from public repositories. Single-cell and single-nucleus RNA-sequencing datasets were obtained from the Gene Expression Omnibus (GEO), including a bladder cancer dataset generated by our group and deposited in GEO (GSE319003). Additional datasets included single-cell RNA-sequencing data from 10 melanoma tumors (GSE148190), 15 liver tumors (GSE125449), and 6 kidney tumors (Broad Single Cell Portal SCP1288; dbGaP phs002065.v1.p1), as well as single-nucleus RNA-sequencing data from 25 bladder tumors (GSE169379), single-cell RNA-sequencing data from 26 breast tumors (GSE176078), lung adenocarcinoma data from 14 patients (GSE123902), and non-small cell lung cancer data from 91 patients (GSE154826). TCGA pan-cancer RNA-sequencing and copy-number alteration data comprising approximately 11,000 samples were obtained from the UCSC Xena Browser (https://xena.ucsc.edu) and cBioPortal for Cancer Genomics (https://www.cbioportal.org). Immunotherapy-response and clinical outcome data from 50 clinical cohorts comprising 5,363 patients were primarily obtained from the Cancer Immunology Data Engine (CIDE) repository (https://cide.ccr.cancer.gov) or from dbGaP through access ID:35035.

### Authors’ contributions

A.R.B., K.E.B., D.Y., and B.L. conceived and designed the study. A.R.B. supervised the research. K.E.B., D.Y. and A.R.B. conducted statistical and bioinformatics analyses of the clinical and multi-omics data. A.R.B., K.E.B. and D.Y. wrote the manuscript. All authors contributed to revision and editing and approved the final manuscript.

## Acknowledgements

This work was supported in part by the Intramural Research Program of the National Institutes of Health (NIH). Contributions by NIH authors were made as part of their official duties as federal employees and are considered works of the United States Government. The findings and conclusions are those of the authors and do not necessarily reflect the views of the NIH or the U.S. Department of Health and Human Services.

