## Supplementary Figures for "APOBEC3G expression marks a TMB-high, T cell–inflamed tumor state and is associated with response to immune checkpoint blockade in multiple cancer cohorts"

### Table of contents

Supplementary Figure 1 (page 2-3)
Supplementary Figure 2 (page 4)
Supplementary Figure 3 (page 5-6)
Supplementary Figure 4 (page 7)
Supplementary Figure 5 (page 8)
Supplementary Figure 6 (page 9-10)

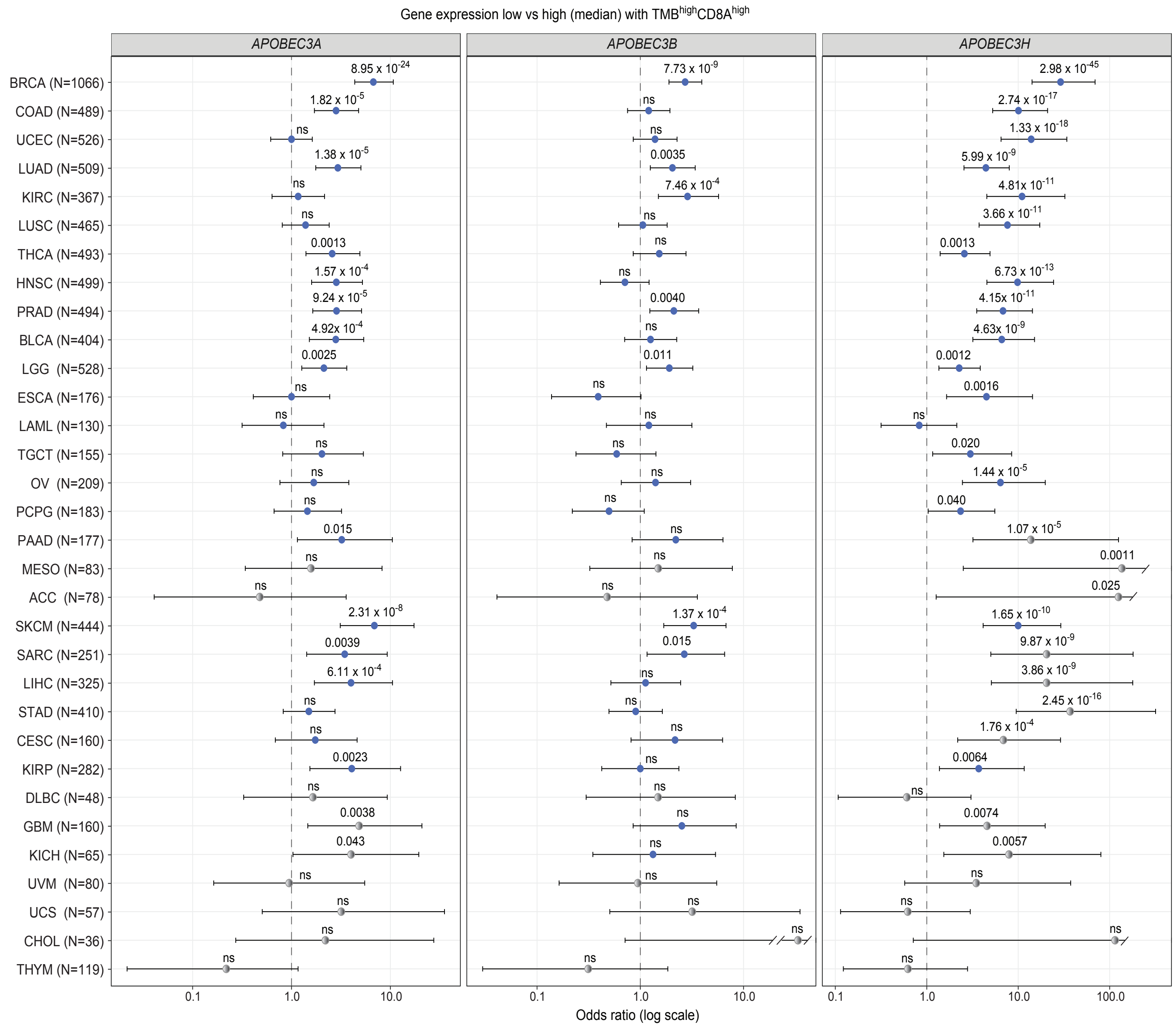


**Supplementary Figure 1. APOBEC3 family members show variable associations with a $TMB^{high}CD8^{high}$ phenotype across cancer types.** Forest plots showing odds ratios (ORs) and 95% confidence intervals (CIs) for the association between expression (high vs. low, median dichotomization) of *APOBEC3A* (left), *APOBEC3B* (middle), and *APOBEC3H* (right) and a $TMB^{high}CD8^{high}$ phenotype across TCGA cancer types. ORs were calculated using Fisher's exact test with Haldane–Anscombe correction to account for zero counts. Cancer types are ordered consistently across panels. Blue points indicate statistically significant associations, while gray points indicate non-significant results. Error bars represent 95% CIs on a log scale. Estimates derived from sparse data (any contingency table cell <5) were retained and should be interpreted with caution. “ns” denotes non-significant associations.

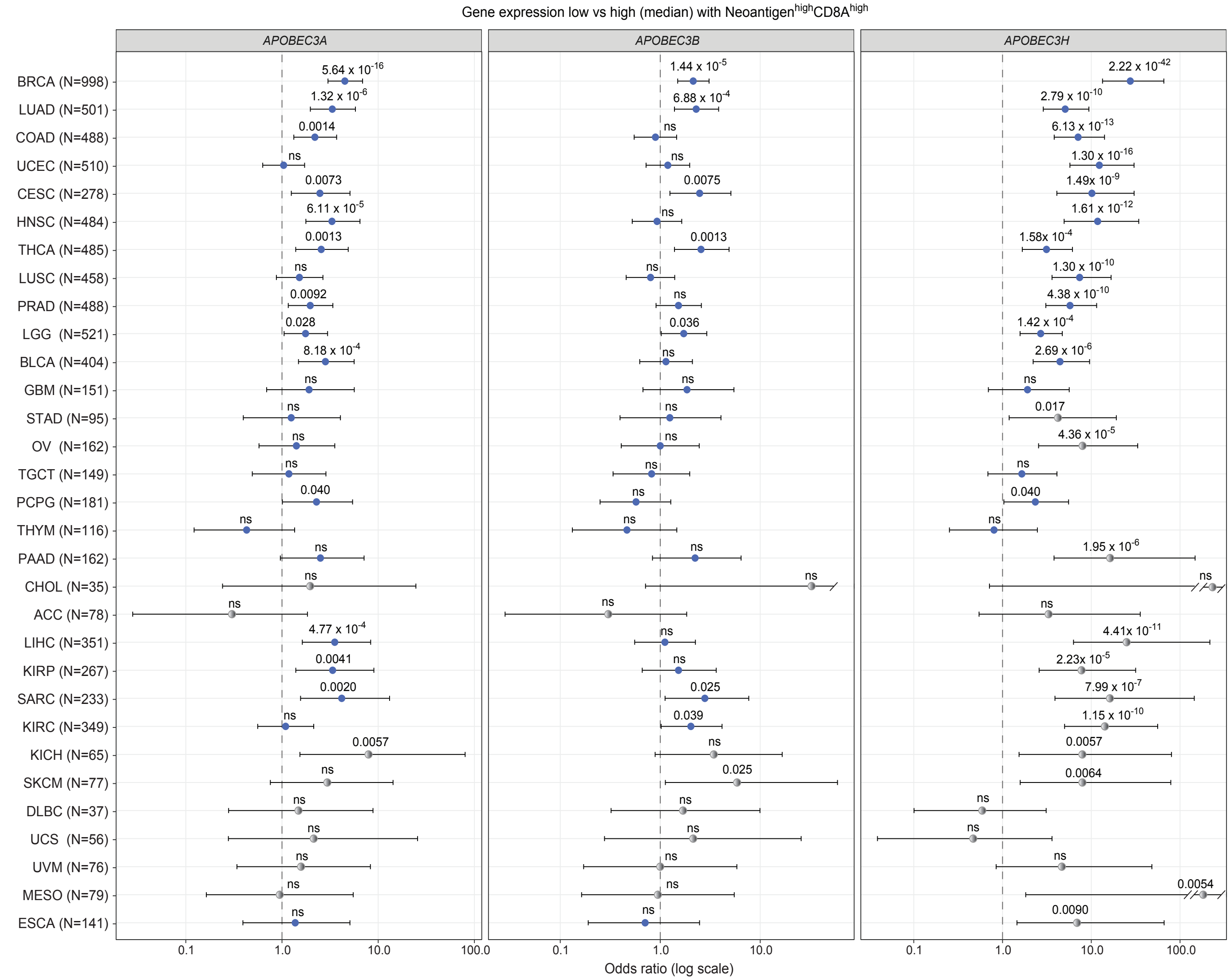


**Supplementary Figure 2. APOBEC3 family members show variable associations with a Neoantigen$^{high}$CD8A$^{high}$ phenotype across cancer types.** The forest plots display the odds ratios (ORs) and 95% confidence intervals (CIs) for the associations between expression (high vs. low, median dichotomization) of *APOBEC3A* (left), *APOBEC3B* (middle), and *APOBEC3H* (right) and a Neoantigen$^{high}$CD8$^{high}$ phenotype across TCGA cancer types. ORs were calculated using Fisher's exact test with the Haldane–Anscombe correction to account for zero counts. The order of cancer types is consistent across all panels. Statistically significant associations are marked with blue points, while non-significant results are shown in gray. The error bars represent the 95% CIs on a logarithmic scale. Estimates derived from sparse data (any contingency table cell < 5) were retained and should be interpreted with caution. "ns" indicates non-significant associations.

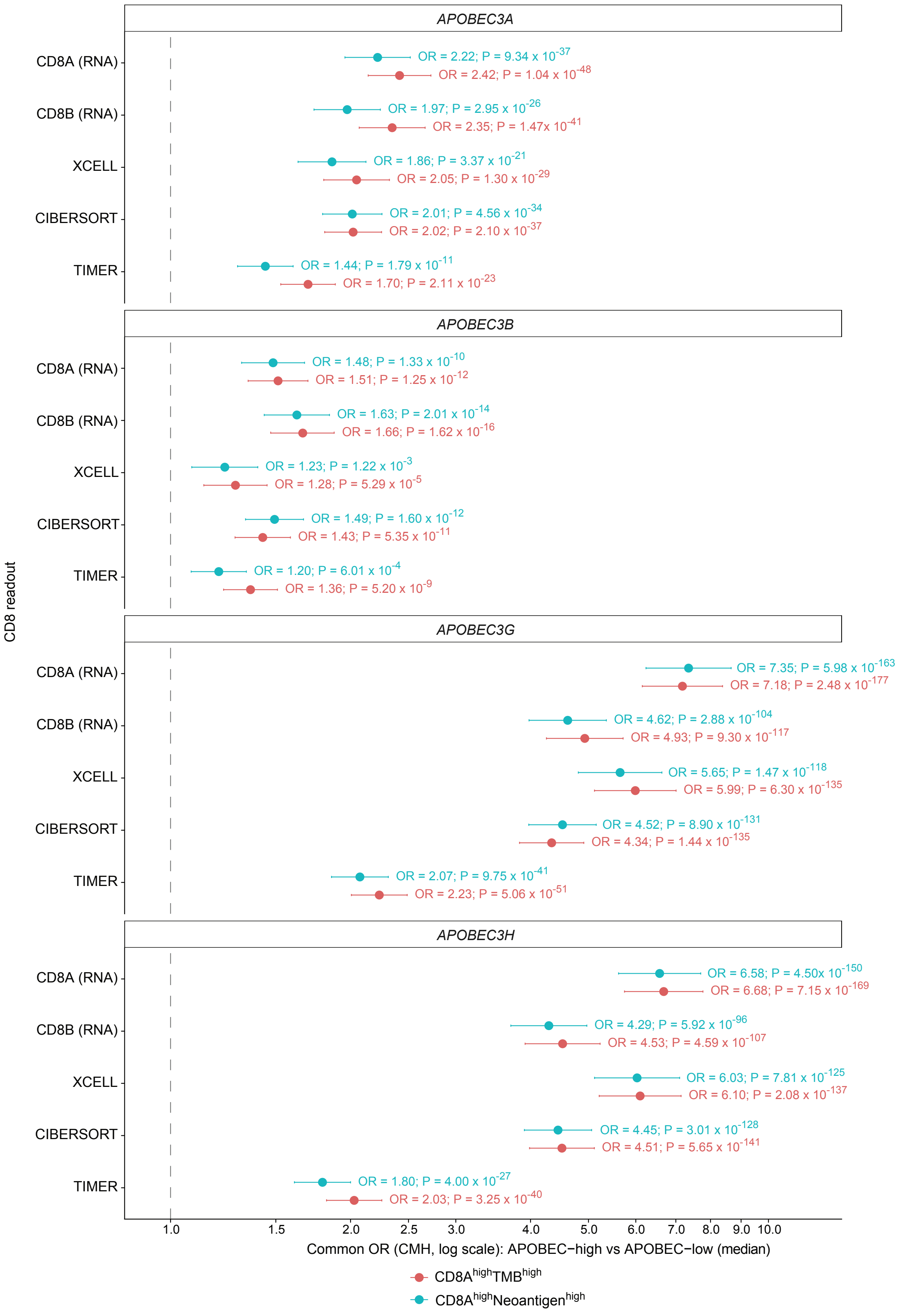
APOBEC3A
CD8A (RNA)
OR = 2.22; P = 9.34 x 10^-37
OR = 2.42; P = 1.04 x 10^-48
CD8B (RNA)
OR = 1.97; P = 2.95 x 10^-26
OR = 2.35; P = 1.47x 10^-41
XCELL
OR = 1.86; P = 3.37 x 10^-21
OR = 2.05; P = 1.30 x 10^-29
CIBERSORT
OR = 2.01; P = 4.56 x 10^-34
OR = 2.02; P = 2.10 x 10^-37
TIMER
OR = 1.44; P = 1.79 x 10^-11
OR = 1.70; P = 2.11 x 10^-23
APOBEC3B
CD8A (RNA)
OR = 1.48; P = 1.33 x 10^-10
OR = 1.51; P = 1.25 x 10^-12
CD8B (RNA)
OR = 1.63; P = 2.01 x 10^-14
OR = 1.66; P = 1.62 x 10^-16
XCELL
OR = 1.23; P = 1.22 x 10^-3
OR = 1.28; P = 5.29 x 10^-5
CIBERSORT
OR = 1.49; P = 1.60 x 10^-12
OR = 1.43; P = 5.35 x 10^-11
TIMER
OR = 1.20; P = 6.01 x 10^-4
OR = 1.36; P = 5.20 x 10^-9
APOBEC3G
CD8A (RNA)
OR = 7.35; P = 5.98 x 10^-163
OR = 7.18; P = 2.48 x 10^-177
CD8B (RNA)
OR = 4.62; P = 2.88 x 10^-104
OR = 4.93; P = 9.30 x 10^-117
XCELL
OR = 5.65; P = 1.47 x 10^-118
OR = 5.99; P = 6.30 x 10^-135
CIBERSORT
OR = 4.52; P = 8.90 x 10^-131
OR = 4.34; P = 1.44 x 10^-135
TIMER
OR = 2.07; P = 9.75 x 10^-41
OR = 2.23; P = 5.06 x 10^-51
APOBEC3H
CD8A (RNA)
OR = 6.58; P = 4.50x 10^-150
OR = 6.68; P = 7.15 x 10^-169
CD8B (RNA)
OR = 4.29; P = 5.92 x 10^-96
OR = 4.53; P = 4.59 x 10^-107
XCELL
OR = 6.03; P = 7.81 x 10^-125
OR = 6.10; P = 2.08 x 10^-137
CIBERSORT
OR = 4.45; P = 3.01 x 10^-128
OR = 4.51; P = 5.65 x 10^-141
TIMER
OR = 1.80; P = 4.00 x 10^-27
OR = 2.03; P = 3.25 x 10^-40
CD8 readout
1.0
1.5
2.0
2.5
3.0
4.0
5.0
6.0
7.0
8.0
9.0
10.0
Common OR (CMH, log scale): APOBEC−high vs APOBEC−low (median)
CD8A^high TMB^high
CD8A^high Neoantigen^high

**Supplementary Figure 3. A*POBEC3G* shows the strongest and most consistent association with double-high phenotypes across CD8$^+$ T cell readouts.** Cochran–Mantel–Haenszel (CMH) summary estimates showing common odds ratios (ORs) and corresponding p-values for the association between *APOBEC3* gene expression (high vs. low, median dichotomization) and double-high phenotypes across multiple CD8$^+$ T cell readouts (CD8A, CD8B, XCELL, CIBERSORT, TIMER). Double-high phenotypes were defined as TMB$^{high}$CD8$^{high}$ (red) or Neoantigen$^{high}$CD8$^{high}$ (cyan). Each panel represents a distinct *APOBEC3* family member (*APOBEC3A, APOBEC3B, APOBEC3G, APOBEC3H*). Points indicate CMH common ORs across cancer types, and horizontal lines represent 95% confidence intervals on a log scale. Estimates were computed by stratifying across cancer types to account for inter-tumor heterogeneity. Across all CD8$^+$ T-cell quantification methods, APOBEC3G showed the strongest and most consistent enrichment for double-high phenotypes among the APOBEC3 family members evaluated, particularly when compared to *APOBEC3A* and *APOBEC3B*.

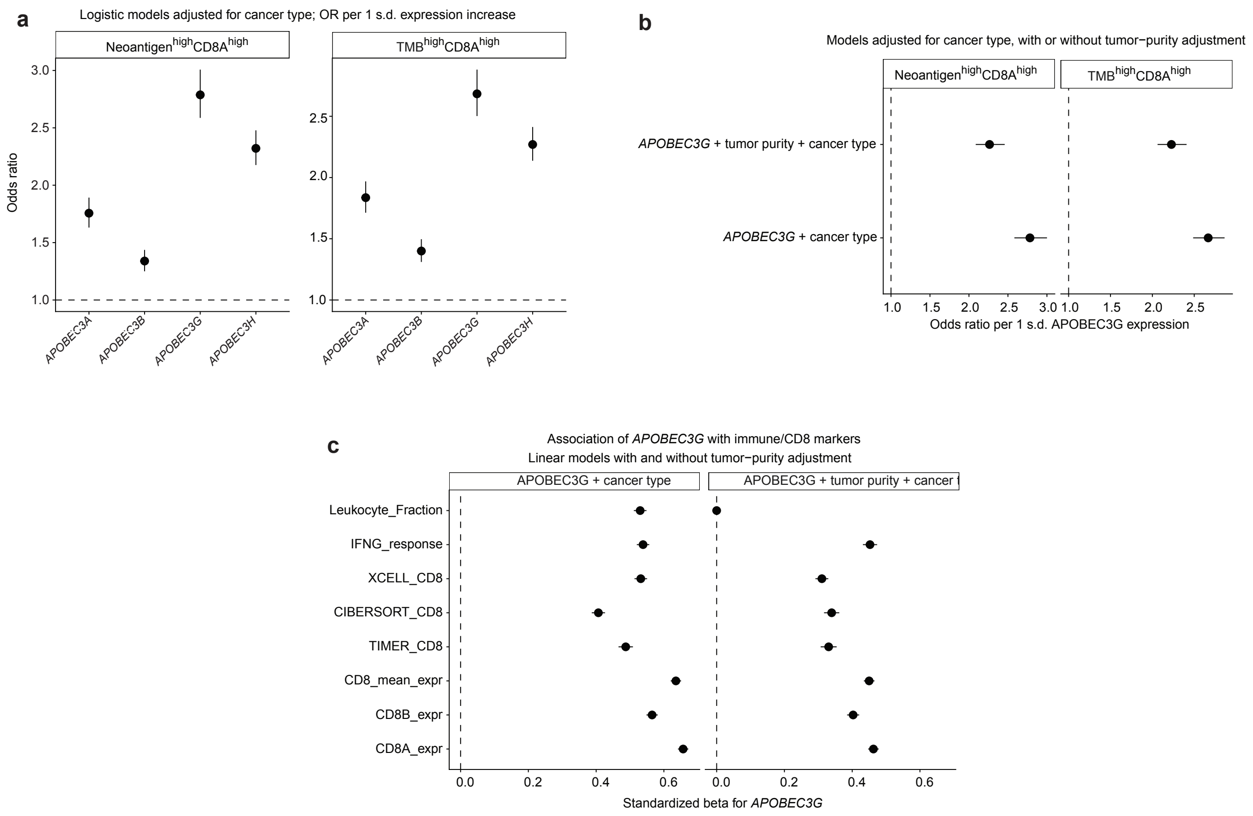


**Supplementary Figure 4. *APOBEC3G* expression is associated with composite tumor mutational burden–immune states after accounting for cancer type and tumor purity.**

**a**, Cancer-type-adjusted logistic regression models evaluating associations between APOBEC3 family members and composite Neoantigen$^{high}$CD8$^{high}$ and TMB$^{high}$CD8$^{high}$ tumor states. Odds ratios are shown per one standard deviation increase in *APOBEC3* expression. *APOBEC3G* showed the strongest association with both composite tumor states among the *APOBEC3* family members tested.

**b**, Sensitivity analysis evaluating the association between *APOBEC3G* expression and composite Neoantigen$^{high}$CD8$^{high}$ and TMB$^{high}$CD8$^{high}$ tumor states in models adjusted for cancer type alone or for both cancer type and tumor purity. *APOBEC3G* remained associated with both composite states after tumor-purity adjustment.

**c**, Linear regression analyses evaluating the association between *APOBEC3G* expression and immune/CD8-related markers in bulk TCGA tumors. Models were adjusted for cancer type alone or for both cancer type and tumor purity. Standardized beta coefficients are shown for *APOBEC3G*. Positive coefficients indicate higher immune-marker levels with higher *APOBEC3G* expression. Error bars indicate 95% confidence intervals.

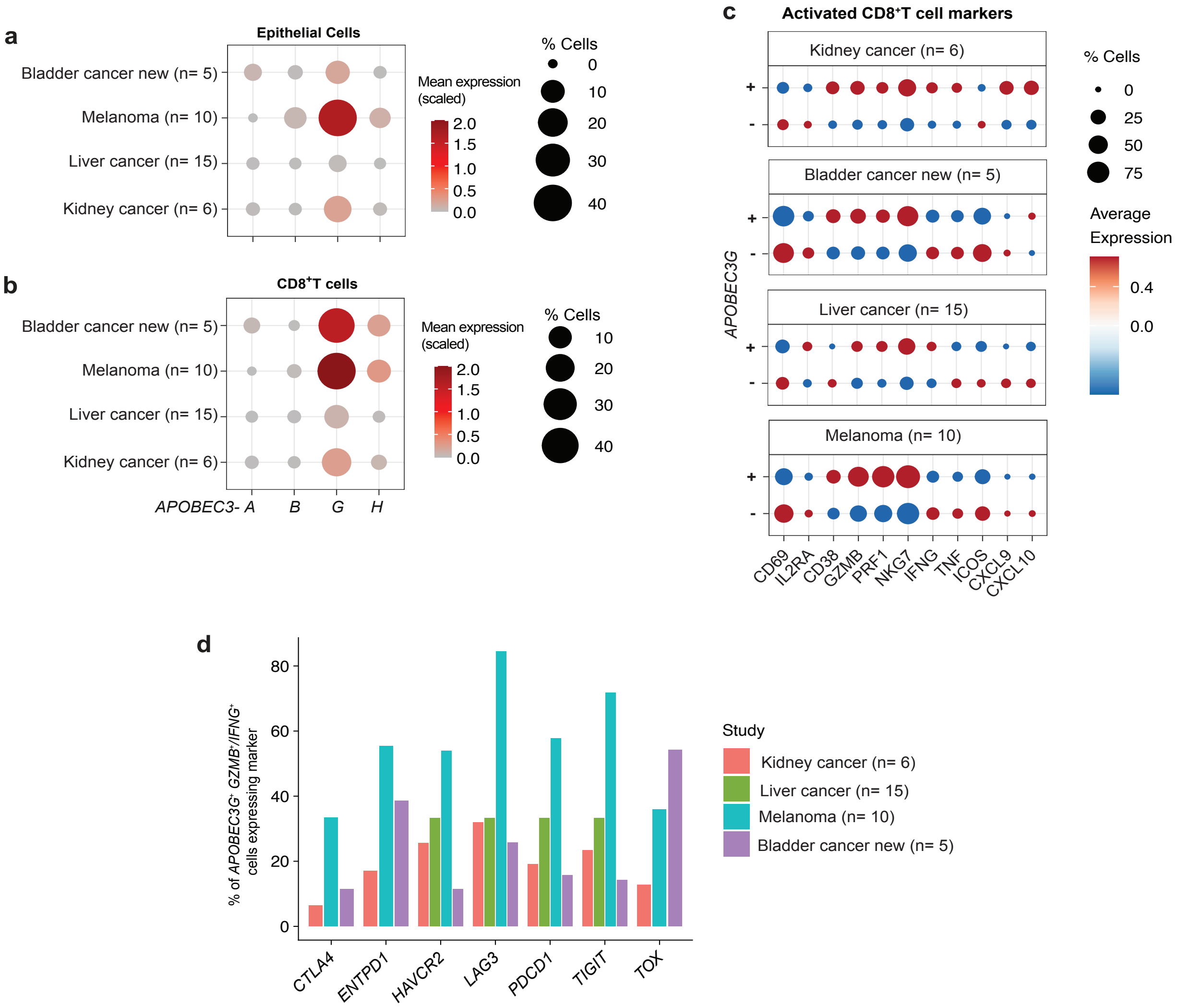


**Supplementary Figure 5. APOBEC3 family expression and activation-marker co-expression in additional independent single-cell tumor datasets.**

**a,b,** Dot plots showing expression of *APOBEC3* family members in malignant epithelial cells (**a**) and CD8+ T cells (**b**) across additional independent single-cell datasets, including a bladder cancer dataset generated by our group and public kidney cancer, liver cancer, and melanoma cohorts. Dot size indicates the percentage of cells expressing each gene, and color indicates scaled mean expression.

**c,** Dot plots comparing activation and effector-marker expression between APOBEC3G-positive and *APOBEC3G*-negative CD8+ T cells within each dataset. Dot size indicates the percentage of cells expressing each marker, and color indicates average expression.

**d,** Bar plot showing the percentage of *APOBEC3G+GZMB+IFNG+ CD8+* T cells co-expressing immune checkpoint or antigen-experience markers across datasets.

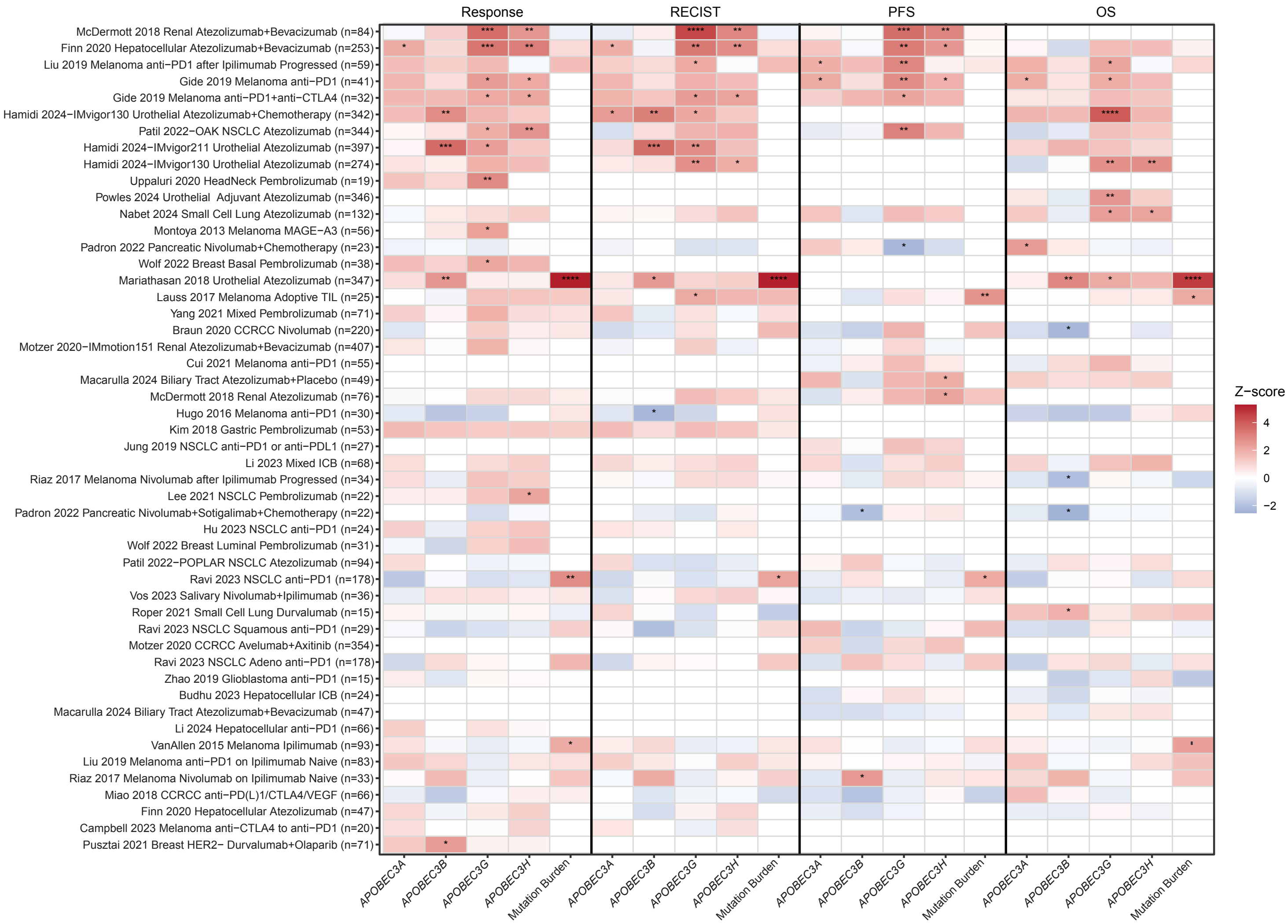


**Supplementary Figure 6. Associations of APOBEC3 family expression and TMB with immunotherapy response and clinical outcomes across multiple cancer cohorts.** Heatmap summarizing associations between expression of APOBEC3 family members (*APOBEC3A, APOBEC3B, APOBEC3G,* and *APOBEC3H)* and tumor mutational burden with clinical outcomes across multiple immunotherapy-treated cohorts from the Cancer Immunology Data Engine (CIDE). Outcomes were grouped into four categories: reported treatment response (Response), RECIST-based response, progression-free survival (PFS), and overall survival (OS). Color intensity represents signed Wald z-statistics derived from logistic regression models for response endpoints and Cox proportional hazards models for survival endpoints, using continuous gene expression values without predefined cutoffs (red, favorable association; blue, unfavorable association). Each row corresponds to an independent cohort, and columns represent genes or biomarkers evaluated within each outcome category. Statistical significance is indicated by asterisks (*$P < 0.05$, **$P < 0.01$, ***$P < 0.001$). Missing values indicate unavailable data for a given cohort or endpoint. Sample sizes indicate the maximum number of evaluable patients across outcome measures. Across cohorts and outcome measures, *APOBEC3G* showed more consistent favorable associations with immunotherapy response and survival than other APOBEC3 family members.
